# Pseudotime analysis reveals exponential trends in DNA methylation aging with mortality associated timescales

**DOI:** 10.1101/2021.11.28.470239

**Authors:** Kalsuda Lapborisuth, Colin Farrell, Matteo Pellegrini

## Abstract

The epigenetic trajectory of DNA methylation profiles has a nonlinear relationship with time, reflecting rapid changes in DNA methylation early in life that progressively slow. In this study, we use pseudotime analysis to determine these trajectories. Unlike epigenetic clocks that constrain the functional form of methylation changes with time, pseudotime analysis orders samples along a path based on similarities in a latent dimension to provide an unbiased trajectory. We show that pseudotime analysis can be applied to DNA methylation in human blood and brain tissue and find that it is highly correlated with the epigenetic states described by the Epigenetic Pacemaker. Moreover, we show that the pseudotime nonlinear trajectory can be modeled using a sum of two exponentials with coefficients that are close to the timescales of human age-associated mortality. Thus, for the first time, we can identify age-associated molecular changes that appear to track the exponential dynamics of mortality risk.

## 1. Introduction

Genome-based studies have provided insights into how DNA regulates biological processes involved in human aging [1,2]. However, DNA is largely invariant during a lifespan of an organism and genetic data provides limited insights on age-associated molecular changes [3]. By contrast, methylation of genomic DNA at cytosine bases is dynamic during our lifespan and is associated with gene regulation and human development [3–6]. Systematic DNA methylation changes proceed predictably with chronological age [7], which has led to the development of various epigenetic aging models that predict age using DNA methylation data [8–13].

Epigenetic aging models, known as epigenetic clocks, can accurately predict chronological age [8,10,13]. The residual term of epigenetic clocks [8,10] has been used to measure biological age acceleration [14], and has been associated with mortality risk [15,16] along with various disease outcomes [17–20]. However, epigenetic clocks are fit by minimizing the difference between observed and predicted ages, which induces linearity between the two measures [8,9,21,22]. Such models are trained to reduce the importance of methylation sites that increase the prediction error but that may be biologically informative, including those that may be associated with nonlinear changes in methylation with age [23,24]. Although some epigenetic clocks attempt to model the age-associated methylation change as some specific nonlinear function of time [10,25], the nonlinear trends captured by these clocks may be biased due to a priori assumption about the functional relationship between epigenetic changes and age.

As an alternative to approaches that predict chronological age, the Epigenetic Pacemaker (EPM) estimates the epigenetic landscape of an individual by modeling changes at each methylation site with respect to time [26]. The EPM approach reveals that DNA methylation patterns of age-associated sites across tissue types change rapidly early in life and then progressively slow as humans age [7]. However, the Epigenetic Pacemaker approach optimizes the parameters of the model by starting with the actual age of each individual, which may introduce some bias. It is therefore of interest to develop an unbiased model of the dynamics of age-associated DNA methylation changes through the human aging process without using chronological age or making a priori assumptions about its relationship with time.

To this end, we propose the use of pseudotime analysis, also called trajectory inference (TI) methods, to model the trajectory of DNA methylation changes over time. Over 70 variations of pseudotime analysis methods have been proposed [27], typically for longitudinal studies of single-cell expression data [27,28]. Pseudotime analysis orders cells based on similarities in their expression patterns and constructs a path along this ordering to assign each cell a value of pseudotime, a latent (unobserved) dimension providing a quantitative measure of the biological progress, such as cell cycle, cell-type differentiation, or cellular activation [27–29].

Generally, pseudotime analyses consist of three main parts: dimensionality reduction, identification of a trajectory, and determination of pseudotime values along the trajectory [30,31]. A cell’s pseudotime is typically the distance along the trajectory between the cell and the origin of the trajectory [31]. Therefore, pseudotime can be understood as an increasing function of chronological time, but not necessarily linear to chronological time [28].

In this study, we demonstrate that pseudotime analysis is applicable for modeling and visualizing the trajectory of DNA methylation changes in blood and brain tissues throughout the human lifespan. Our results show that there is a significant correlation between the methylation pseudotime and the non-linear dynamics of epigenetic landscapes captured by the Epigenetic Pacemaker Model [26]. Moreover, the methylation pseudotime trajectory is well described by the sum of two exponential trends across a lifespan and suggests that the timescale of the methylation pseudotime may be related to the human population mortality rate.

## 2. Materials and Methods

### 2.1 Methylation Array Processing

Illumina Infinium HumanMethylation450 BeadChip microarray data from whole blood and brain samples in the Gene Expression Omnibus (GEO) [32] with more than 50 samples that do not have missing methylation beadchip array intensity data (IDAT) files, repeated measurements of the same samples, and data from cultured cells or cancerous tissues [24] were selected and processed using minfi [33] (v1.34.0). To mitigate variations between different experiments, whole blood samples were selected if their median methylation probe intensity was greater than 10.5 and the difference between the observed and expected median unmethylation probe intensity was less than 0.4 [24]. We refer the reader to [24] for further processing and selection details for whole blood array samples.

### 2.2 Pseudotime analysis

The pseudotime trajectory was inferred by, first, reducing the dimensionality of the input methylation data matrix with principal component analysis using prcomp in stats R package [34]. Then, we use mclust, an R package for Gaussian Mixture Modelling (GMM) [35], to cluster our samples in the low-dimensional space. The number of clusters and the covariance parameterization were selected based on the Bayesian Information Criterion (BIC) as implemented in mclust [35]. Lastly, the pseudotime value for each training sample was assigned by Slingshot [36]. Slingshot first constructs a minimum spanning tree (MST) on the cluster centers, then applies a simultaneous principal curves method to obtain smooth trajectories of the MST, and finally assigns the pseudotime value for each sample according to the distance from the start of the trajectory to where the sample is orthogonally projected onto the curve [36]. The inferred trajectory was used to predict pseudotime values for the validation and test data that were projected onto the same principal components as the training set. The pseudotime analysis was conducted with a fixed random seed.

### 2.3 Epigenetic Pacemaker (EPM)

The Epigenetic Pacemaker (EPM) measures the epigenetic state of each sample by modeling methylation levels at each CpG site as a linear function of the epigenetic state (i.e. epigenetic age) of an individual [26]. Specifically, a methylation value ***m***_***ij***_, where ***i*** denotes the methylation site and ***j*** the sample, is modeled as

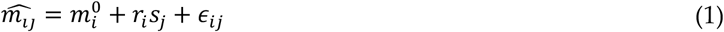

, where

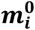 is the initial methylation value,

***r***_***i***_ is the rate of change,

***s***_***j***_ is the epigenetic state, and

***ϵ***_***ij***_ is the normally distributed error term.

The EPM starts with an input matrix of methylation values and uses a fast conditional expectation-maximization algorithm to output the optimal values of 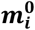, ***r***_***i***_, and ***s***_***j***_ by minimizing the error between the predicted and the input methylation values across all methylation sites. Chronological age was used as the initial guess for epigenetic states for each sample [26].

### 2.4 Determining the functional form of methylation pseudotime

To determine the functional form that best describes relationships between methylation pseudotime values and chronological age as well as epigenetic age, we find the best fitting trend line with five functional forms,

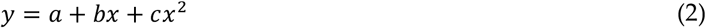

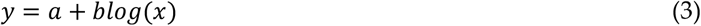

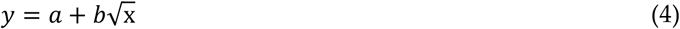

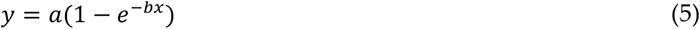

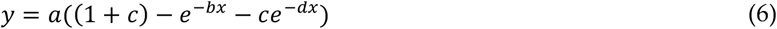

, where

***y*** is methylation pseudotime value

***x*** is chronological age, and

***a, b, c***, and ***d*** are coefficients.

We denote the functions as quadratic (2), logarithmic (3), square root (4), exponential (5), and the sum of two exponentials (6) respectively. Our best-fit criteria include Akaike information criterion (AIC), R-squared value (R2), and Root Mean Square Error (RMSE). The AIC value is calculated using the stats R package [34]. We also use the package to compute the correlation between pseudotime values and epigenetic ages using Pearson’s correlation coefficient (PCC) [34].

### 2.5 Analysis environment

Methylation data processing, site selection, epigenetic state detection was carried out using Python (v.3.9.6) in JupyterNotebook [37] using numpy [38], scikit-learn [39], EpigeneticPacemaker [26], Pandas [40], and Joblib [41] packages. Trajectory inference and additional analysis were carried out with R (v.4.1.1) [34] in RStudio [42]. The mclust [35], slingshot [36], RColorBrewer [43], ggplot2 [44], viridis [45], splitTools [46], and nlsr [47] packages were used.

## 3. Results

### 3.1 Methylation Pseudotime is nonlinear across the lifespan

#### 3.1.1 Methylation Pseudotime of Whole Blood Tissue

Four aggregate datasets composed of Illumina 450k array data from blood samples [48–51] (n=1605, age=0-94 years) were used for inferring a pseudotime trajectory. The samples were stratified and split according to age for pseudotime trajectory training (n=1283) and validation (n=322). For all samples, methylation values at each site were quantile normalized by probe type [10] using the median methylation value of the training blood samples to mitigate batch effects. For training, we selected age-associated methylation sites with the absolute value of the Pearson correlation coefficient (PCC) larger than 0.4 between age and methylation values of training samples (n=8510). Pseudotime analysis clusters a low-dimensional representation (PC=2) of training samples and constructs a pseudotime trajectory across the cluster centers (**Figure 1a**). The inferred trajectory was used to assign pseudotime values of each training and validation sample projected on the same low-dimensional space. We observe that the pseudotime value of each data point increases along the trajectory in the latent space for both training (**Figure A1a**) and validation set (**Figure A1b**). The trained model performed well on the training (R^2^=1.00, RMSE=0.0133) and validation (R^2^=0.996, RMSE=0.236) datasets. **Figure A2a and A2b** show plots of the assigned pseudotime values against chronological age of the training and validation set respectively.

**Figure 1.**
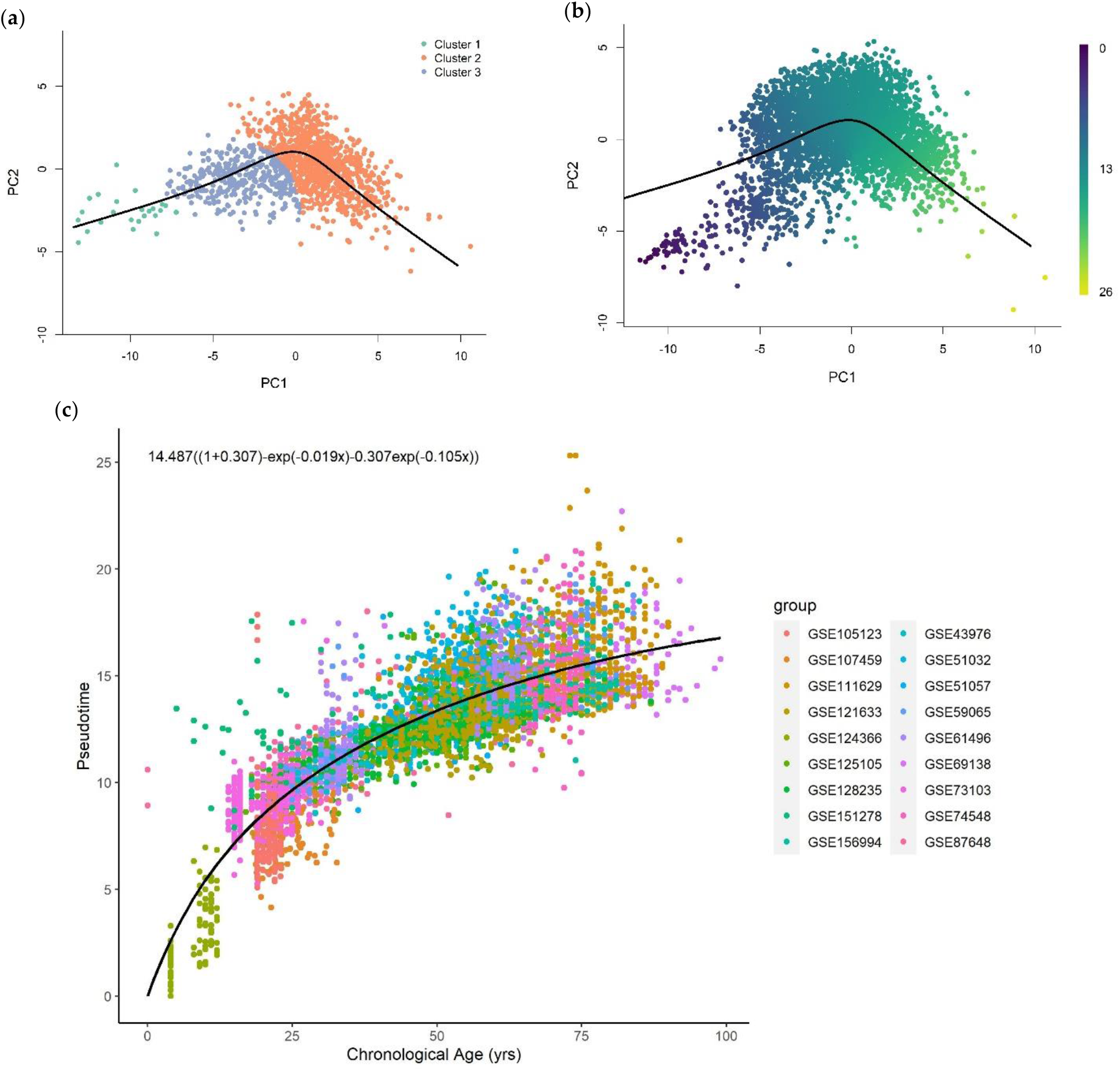
(**a**) The pseudotime trajectory inferred from clusters of training methylation array data of whole blood tissue in low dimensional space (PC=2); (**b**) Methylation array data of whole blood tissue were projected onto low-dimensional space used for training. Each data point is colored according to the methylation pseudotime predicted with the pseudotime trajectory shown; (**c**) Methylation pseudotime of whole blood tissue samples.

The trajectory was used to predict pseudotime of a large aggregate data set composed of Illumina 450k array data [18,52–68] (n=4519, age=0-99 years). The pseudotime value was predicted for each test sample projected onto the reduced-dimensional space used to construct the trajectory (**Figure 1b**). **Figure 1c** shows that the relationship between the predicted pseudotime and chronological age is characterized by a positive nonlinear trend with a rapid change in young samples that slows for older samples.

#### 3.1.2 Methylation Pseudotime of Brain Tissue

A dataset of brain methylation array samples [6] (n=675, age=0-97 years) is used for both trajectory fitting and pseudotime prediction. The samples were stratified and randomly split according to age for pseudotime trajectory training (n=138), validation (n=70), and testing (n=467). Age-associated methylation sites were selected such that the absolute value of PCC between age and methylation values of training samples at each site is larger than 0.70 (n=9446). The trajectory was inferred using a low dimensional representation (PC=2) of training samples at age-associated sites (**Figure 2a**). We then use the trajectory to assign pseudotime values to training and validation samples projected onto the low-dimensional space. The pseudotime value for both training and validation samples increases along the trajectory as shown in **Figure A3a** and **Figure A3b** respectively. The trajectory was able to predict the pseudotime value well on the training (R^2^=1.00, RMSE=0.00939) and validation (R^2^=0.980, RMSE=1.108) datasets. The plots of the assigned pseudotime values against chronological age are shown in **Figure A4a** for the training set and **Figure A4b** for the validation set.

**Figure 2.**
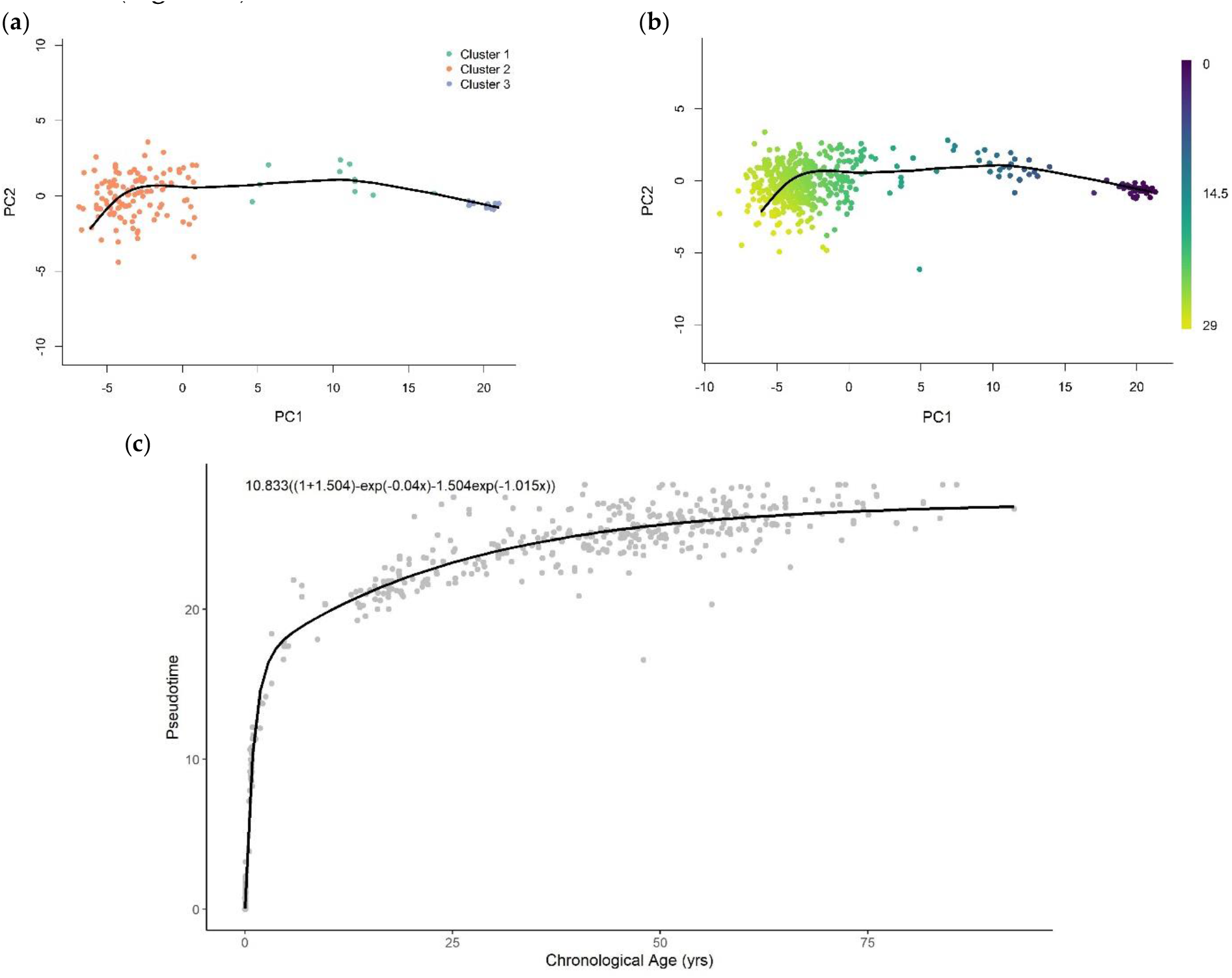
(**a**) The pseudotime trajectory inferred from clusters of training methylation array data of brain tissue in low dimensional space (PC=2); (**b**) Methylation array data of brain tissue were projected onto low-dimensional space used for training. Each data point is colored according to the methylation pseudotime predicted with the pseudotime trajectory shown; (**c**) Methylation pseudotime of brain tissue samples.

The representations of samples in the same reduced-dimensional space as model training were used for pseudotime value prediction of the test set (**Figure 2b**). Similar to pseudotime of blood samples, the predicted pseudotime of brain samples rapidly increases in young samples and decelerates as age increases (**Figure 2c**).

### 3.2 Methylation pseudotime is linearly correlated to epigenetic states estimated by the EPM

Since it is shown that the change of epigenetic state across the lifespan is characterized by a similar nonlinear trend to that of methylation pseudotime [7], we investigated the relationship between the two values. We fit cross-validated (cv=5) EPM [26] models to each of the blood and brain tissue training methylation array data at the same selected age-associated sites as previously described for pseudotime analysis. We then use the fitted EPM model to predict the epigenetic state for samples in the test set. We test the correlation between pseudotime and the estimated epigenetic state using the Pearson Correlation Coefficient (PCC) and fit a linear function using the stats R package [34].

**Figure 3** and **Figure 4** illustrate a clear linear relationship between the pseudotime and the estimated epigenetic state of blood and brain samples respectively. The statistically significant PCC values (r_blood_=0.972, P_blood_<0.001; r_brain_=0.997, P_brain_<0.001) further point towards a strong linear correlation between the two values for both tissue types.

**Figure 3.**
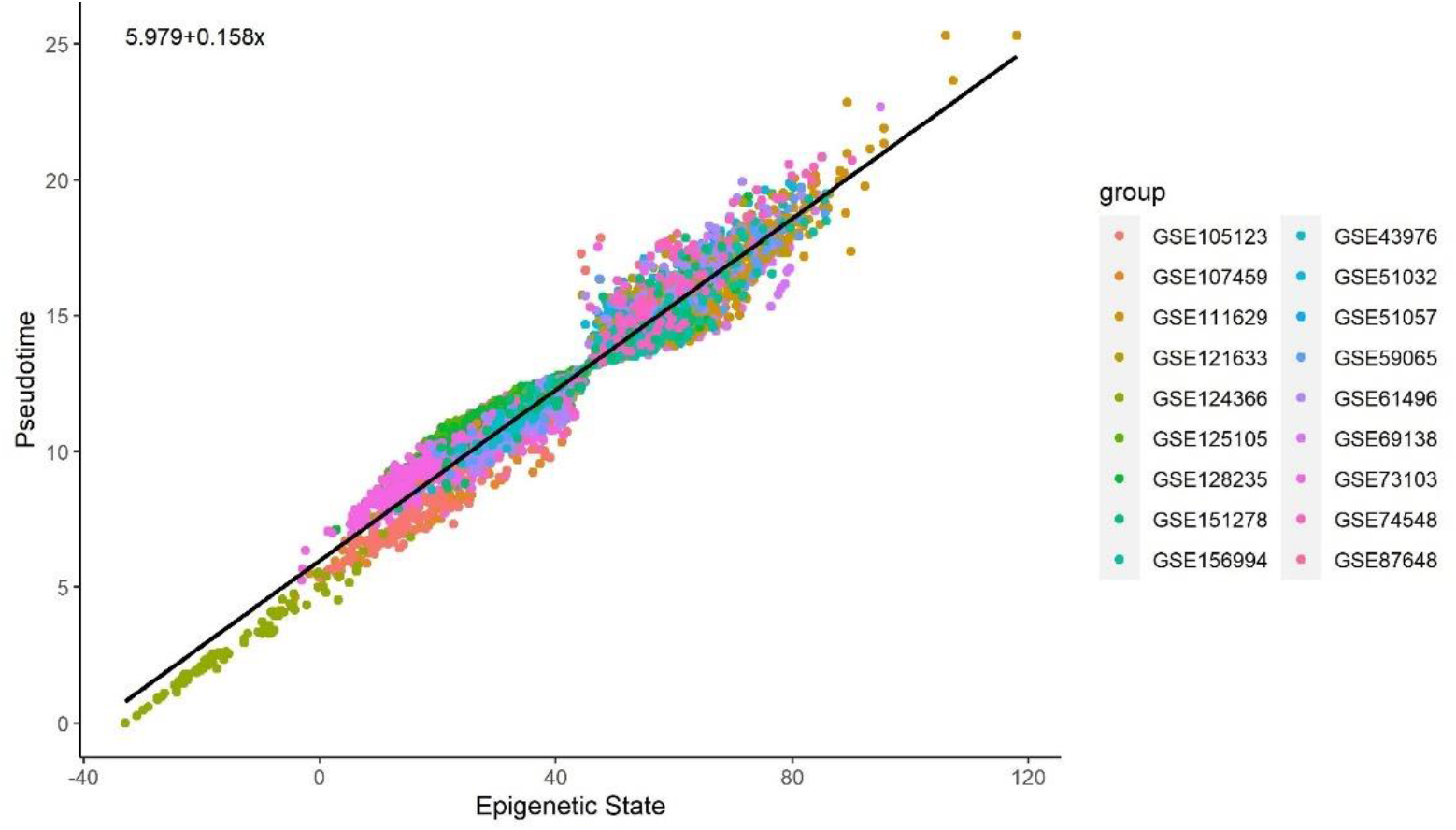
Methylation pseudotime vs Epigenetic State of whole blood tissue samples

**Figure 4.**
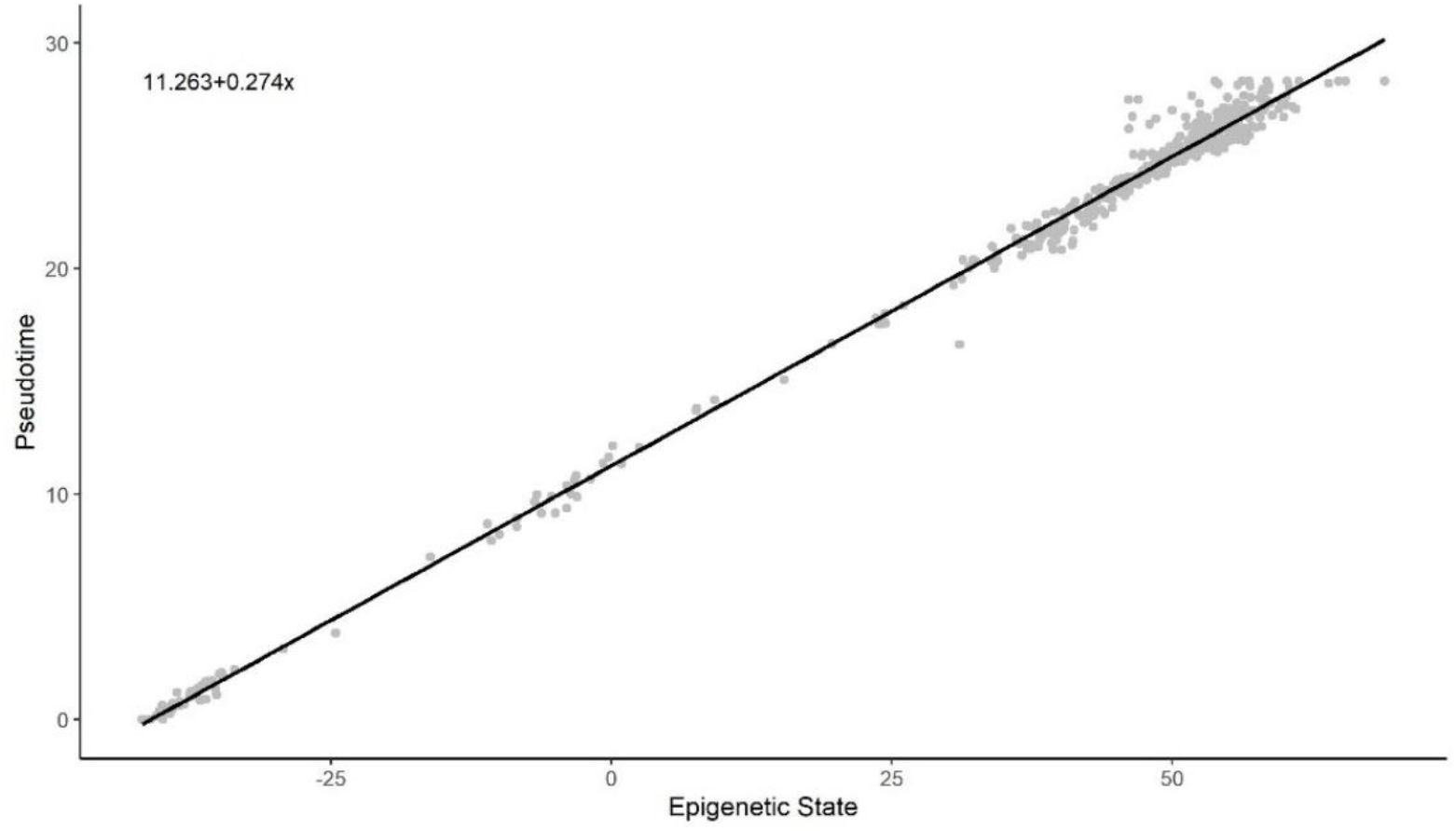
Methylation pseudotime vs Epigenetic State of brain tissue samples

### 3.3 Sum of Two Exponentials Best Describe Methylation Pseudotime Across Lifespan

Since the relationship between pseudotime and chronological age in both blood and brain tissue are similarly characterized by an initial rapid change that slows as the sample age increases, we sought a function that best describes this nonlinear trend by considering the AIC, RMSE, and R2 value, as measures of fit. The coefficients for the quadratic (**2**), logarithmic (**3**), square root (**4**), exponential (**5**), and the sum of two exponential (**6**) functions were estimated with the stats R package [34].

#### 3.3.1 Functional form of Methylation Pseudotime of Whole Blood Tissue

All functional forms can describe the general trend of pseudotime across the entire lifespan (**Figure A5**). The logarithmic, exponential, and sum of two exponential trend lines generate improved fits to the methylation pseudotime of young samples. Amongst all functions, the sum of two exponentials had the highest R-squared value with the lowest AIC and RMSE values (**Table 1**). This suggests that the sum of two exponentials approximates methylation pseudotime as a function of chronological age more accurately than all other functional forms. Therefore, the sum of two exponentials best describes the change of methylation pseudotime of whole blood samples across the lifespan (**Figure 1C**).

**Table 1.** Fit metrics of logarithmic, square root, exponential, and sum of two exponential functions to methylation pseudotime of whole blood samples

| Functions with estimated coefficients | AIC | RMSE | R<sup>2</sup> |
| --- | --- | --- | --- |
| $y = -5.486 + 4.825\log(x)$ | 18258.27 | 1.773 | 0.671 |
| $y = 1.23 + 1.686\sqrt{x}$ | 17991.60 | 1.722 | 0.690 |
| $y = 3.56 + 0.277x - 0.002x^2$ | 17896.36 | 1.704 | 0.697 |
| $y = 16.531(1 - e^{-0.035x})$ | 17913.38 | 1.708 | 0.695 |
| $y = 14.487((1 + 0.307) - e^{-0.019x} - 0.307e^{-0.105x})$ | 17826.79 | 1.690 | 0.700 |

#### 3.3.2 Functional form of Methylation Pseudotime of Brain Tissue

Only the logarithmic and sum of two exponential functions describe the data well across the entire lifespan (**Figure A6**). The sum of two exponentials has the highest R-squared value with the lowest AIC and RMSE values for all functions (**Table 2**). This indicates that the sum of two exponentials approximates methylation pseudotime as a function of chronological age more accurately than all other functional forms. Therefore, the sum of two exponentials best describes the change of methylation pseudotime of brain tissue samples across the lifespan (**Figure 2C**).

**Table 2.**
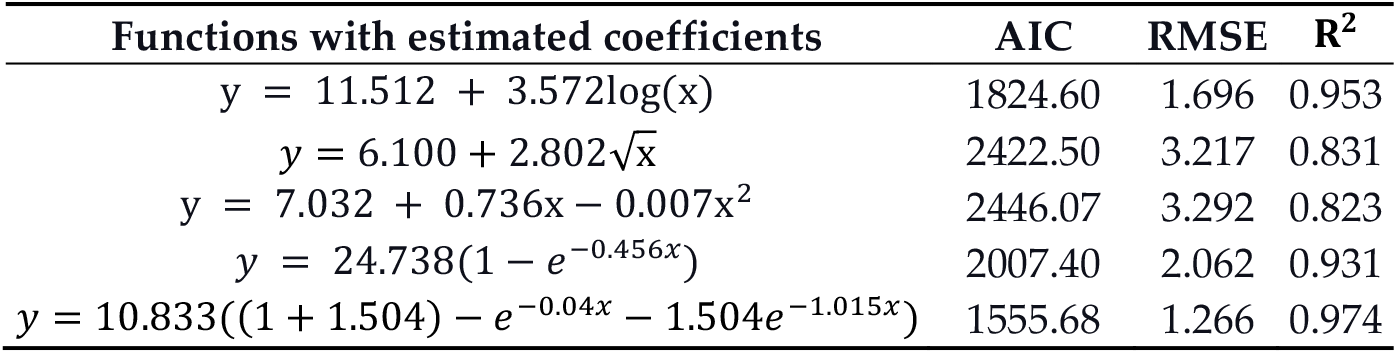
Fit metrics of logarithmic, square root, exponential, and sum of two exponential functions to test samples of methylation pseudotime of brain samples

| Functions with estimated coefficients | AIC | RMSE | R<sup>2</sup> |
| --- | --- | --- | --- |
| $y = 11.512 + 3.572\log(x)$ | 1824.60 | 1.696 | 0.953 |
| $y = 6.100 + 2.802\sqrt{x}$ | 2422.50 | 3.217 | 0.831 |
| $y = 7.032 + 0.736x - 0.007x^2$ | 2446.07 | 3.292 | 0.823 |
| $y = 24.738(1 - e^{-0.456x})$ | 2007.40 | 2.062 | 0.931 |
| $y = 10.833((1 + 1.504) - e^{-0.04x} - 1.504e^{-1.015x})$ | 1555.68 | 1.266 | 0.974 |

## 4. Discussion

Several studies proposed epigenetic aging models, or the epigenetic clocks, that predict the age of an individual based on the changes of DNA methylation profile over time [8,9,13]. The residuals of these epigenetic aging models are used as aging and health biomarkers [14–20]. Epigenetic clocks are generally formulated to discount the importance of methylation sites that increase the age prediction error, including those associated with a nonlinear change in methylation with age [23,24]. This results in the modeled methylation change being linear with age [8,9,21,22]. Some epigenetic clocks account for the nonlinearity by modeling epigenetic aging as a nonlinear function of chronological age [10,25]. Nonetheless, the methylation aging trend captured by epigenetic clocks may be biased due to a priori assumption about the functional form underlying the relationship between methylation and chronological age.

The Epigenetic Pacemaker (EPM) provides a different approach to model age-associated methylation changes by directly modeling the methylation matrix as a time-dependent entity. The EPM shows that DNA methylation changes at age-associated sites are nonlinear with time [7]. However, the EPM paradigm used the known age of each individual as the starting value of epigenetic age as its initial condition for optimization, which may generate some bias in the estimation. It is therefore of interest to also develop a model of the dynamics of DNA methylation changes through the human aging process that is completely independent of the underlying relationship between time-dependent methylation changes with chronological age.

To accomplish this, we used pseudotime analysis to learn the trajectory of DNA methylation through the human aging process and investigate the correlation between methylation pseudotime and the epigenetic state estimated by the EPM. By applying the pseudotime analysis to DNA methylation data taken from a dataset of brain samples and an aggregated dataset of blood samples, we observe consistent trends for both datasets. Across both tissue types, the relationship between chronological age and the pseudotime values is characterized by a nonlinear trend with rapid changes in young samples that slow as sample ages increase.

We observe that the methylation pseudotime and the epigenetic states estimated by the EPM have a statistically significant linear correlation. This suggests that pseudotime is closely tracking epigenetic states. Since methylation pseudotime values were estimated without using chronological age, its correlation to the epigenetic state also provides biological support for the nonlinear epigenetic aging trends first captured by the EPM.

To quantitatively understand the timescales over which DNA methylation is changing across a lifespan we fit the pseudotime trajectories to various analytical functions. We find that the observed nonlinear trends are best described by the sum of two exponentials. Since we are able to detect such trends across different tissue types, the sum of two exponential functions may provide a consistent description of the human epigenetic aging trajectories. The sum of two exponential trends can be interpreted as describing two independent exponential processes that occur simultaneously, one of which is faster than the other. Age-related human mortality is characterized by the Gompertz equation [69], *μ(x)* = *αe*^*βx*^, such that *μ(x)* is the mortality rate at age *x* and *β* describes how fast the mortality rate increases with time. *β* for the human population is found to be 0.08, which results in a doubling of mortality risk every 8.5 years [70,71]. For the whole blood dataset, we find that one of the two coefficients is 0.1, which results in a doubling time of 6.93 years. This coefficient is remarkably close to 0.08, the Gompertz coefficient. This suggests that the timescale of epigenetic changes in the blood are similar to those observed in mortality risk, both of which have a doubling time around ten years. By contrast, the slower time scale of the brain data is slightly slower than the Gompertz beta, at 0.04 instead of 0.08. The beta of the brain data results in a doubling time of 17.33 years instead of 8.5 years. This may indicate that brains age epigenetically at a slower rate than blood, although even the brain’s exponential timescale is not far off from the Gompertz one. Moreover, the faster brain timescale we observe in our double exponential fit may reflect the very rapid changes that occur during development in the first few years of life.

What is the implication of the observation that DNA methylation patterns change exponentially at timescales similar to those observed in human mortality studies? One hypothesis is that the epigenome is set early in development, and then as we age the methylation patterns randomly drift and become more diverse due to epimutations and changes in immune cell type composition. This drift is characterized by an exponential process, and therefore is asymptotically approaching a level of maximum diversity or entropy, at which point the organism has the maximum risk of dying. In other words, as we age, we are exponentially losing the epigenetic specification of our cells, and this process is tracking the exponentially increasing risk of mortality. In support of this hypothesis, we show that the timescales of these molecular changes are similar to the timescales of mortality risk.

In conclusion, we demonstrate that pseudotime analysis can be applied to DNA methylation data to model the trajectory of its changes over time without relying on a priori knowledge of the functional form of methylation changes with time. The strong correlation between methylation pseudotime and epigenetic states suggests that both approaches are capturing the nonlinear trends in the dynamics of epigenetics during a lifespan. Moreover, we provide evidence that the timescale for these processes is intriguingly similar to the timescale of mortality risk, further buttressing the hypothesis that aging is at least in part driven by epigenetic drift.

## Supplementary Materials

Analysis code can be found at https://github.com/Kal-Lap/PseudotimeMethylation.

## Author Contributions

K.L., methodology, software, formal analysis, visualization, and writing-original draft preparation; C.F., data curation and writing-review and editing; M.P., conceptualization and writing-review and editing. All authors have read and agreed to the published version of the manuscript.

## Institutional Review Board Statement

Not applicable.

## Informed Consent Statement

Not applicable.

## Data Availability Statement

Publicly available datasets from the Gene Expression Omnibus (GEO) repository (https://www.ncbi.nlm.nih.gov/geo/) were used for analysis in this study. The datasets can be accessed with the following accession number: GSE42861, GSE43976, GSE51032, GSE51057, GSE59065, GSE61496, GSE69138, GSE73103, GSE74193, GSE74548, GSE87571, GSE87648, GSE97362, GSE105123, GSE107459, GSE111629, GSE121633, GSE124366, GSE125105, GSE128064, GSE128235, GSE151278, GSE156994.

## Appendix

**Figure A1.**
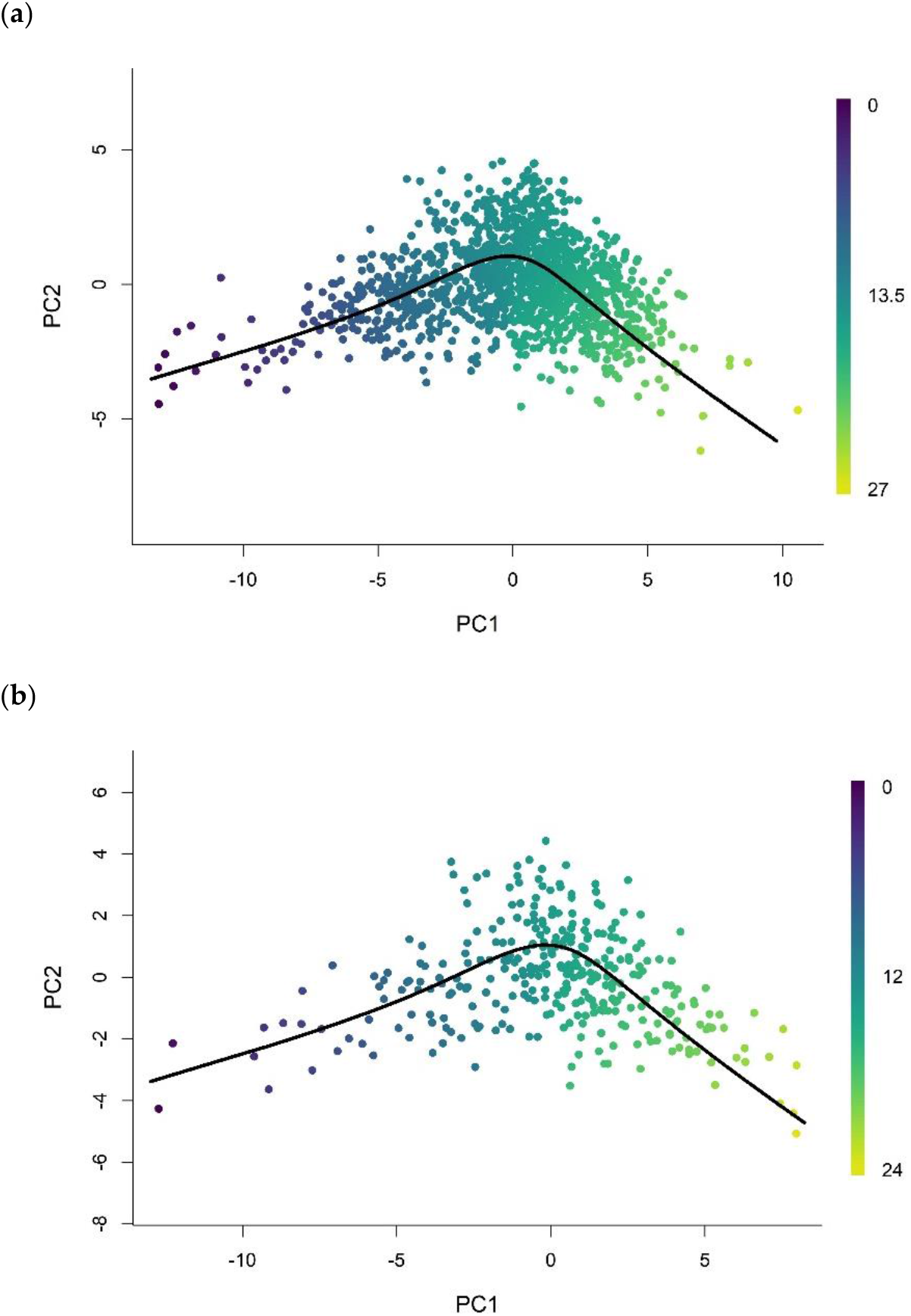
Methylation array data of whole blood tissue in (**A**) train and (**B**) validation set were projected onto low-dimensional space (PC=2) used for training. Each data point is colored according to the methylation pseudotime predicted with the pseudotime trajectory shown.

**Figure A2.**
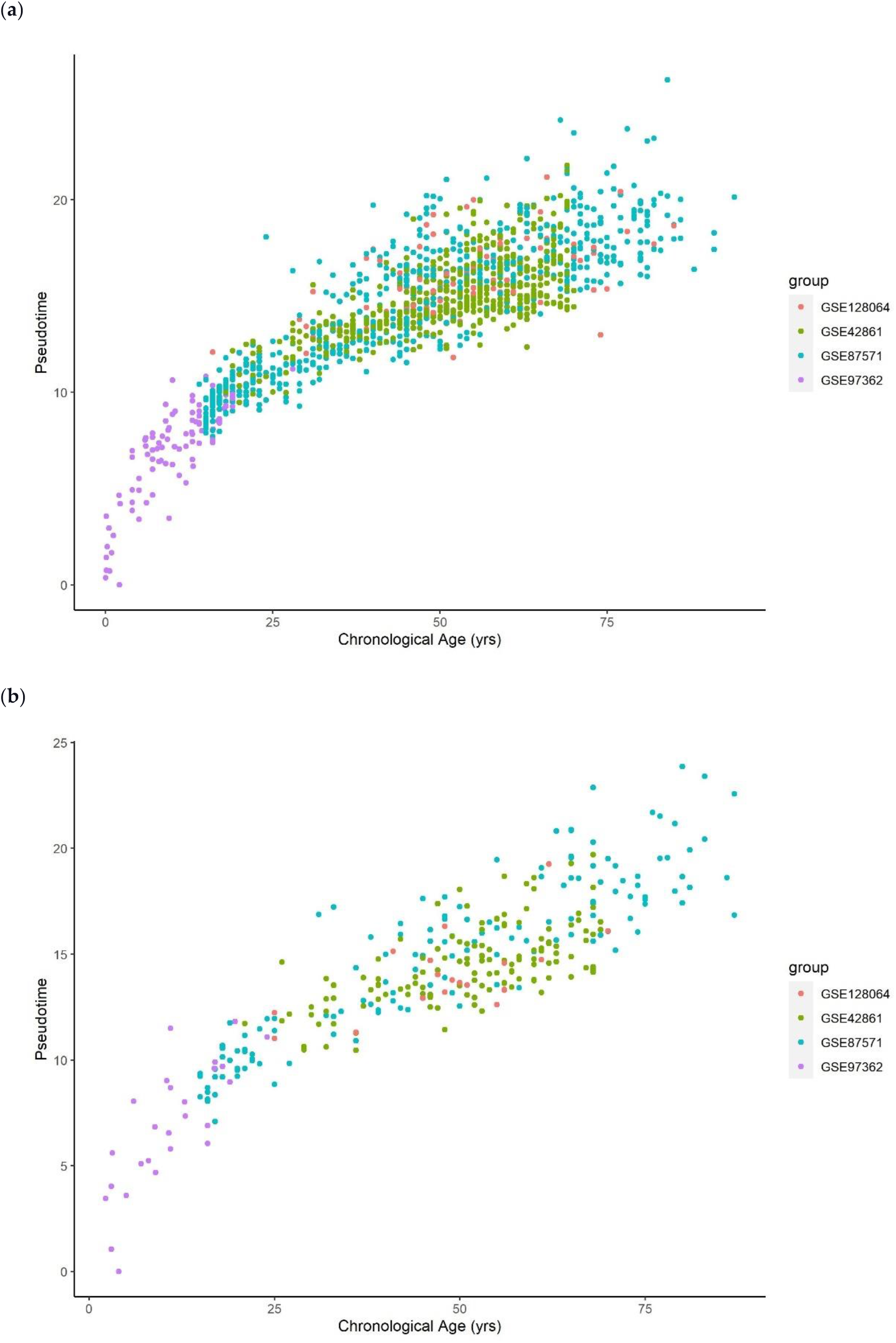
Methylation pseudotime of whole blood tissue samples in (**A**) train and (**B**) validation set.

**Figure A3.**
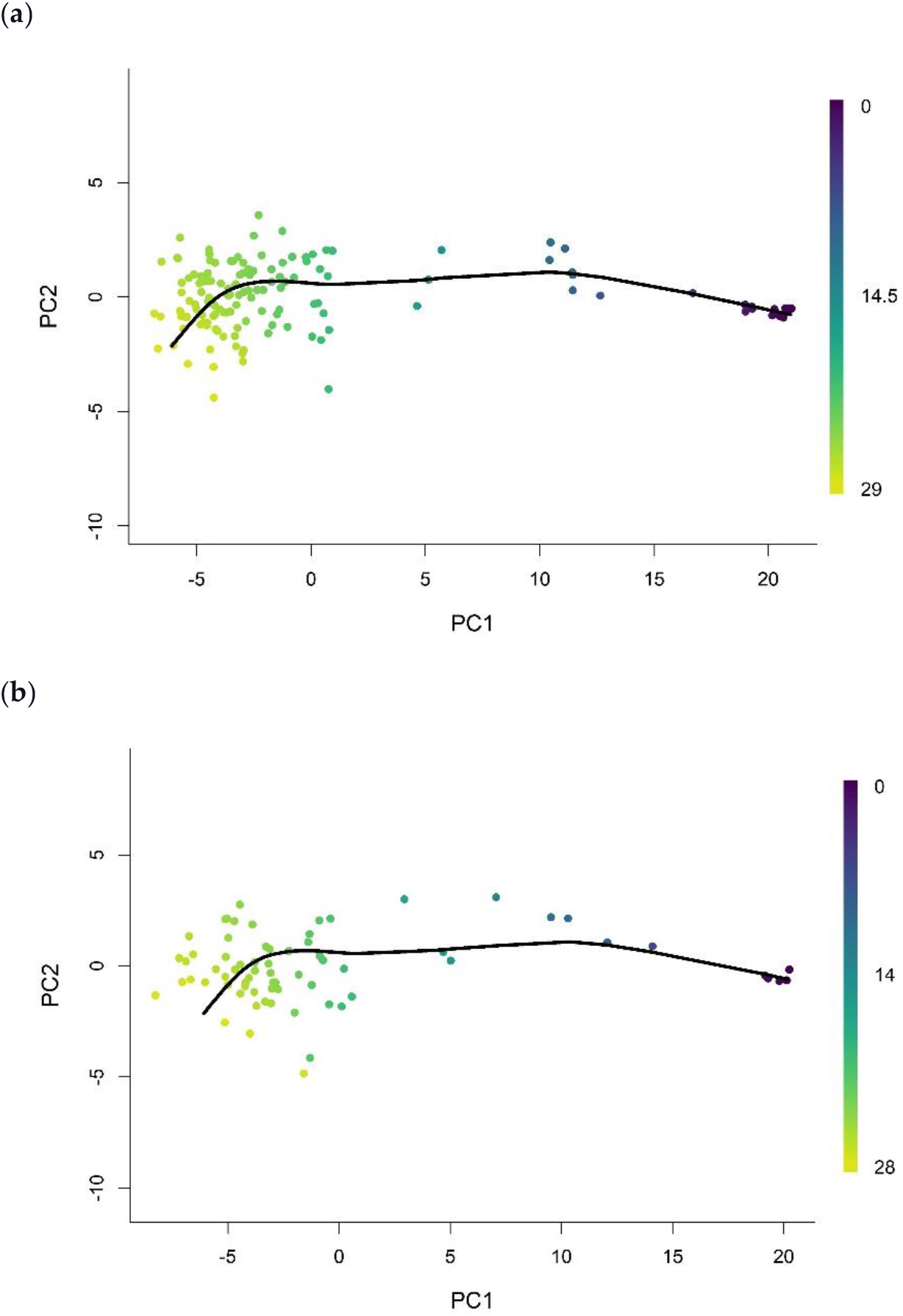
Methylation array data of brain tissue in (**A**) train and (**B**) validation set were projected onto low-dimensional space (PC=2) used for training. Each data point is colored according to the methylation pseudotime predicted with the pseudotime trajectory shown.

**Figure A4.**
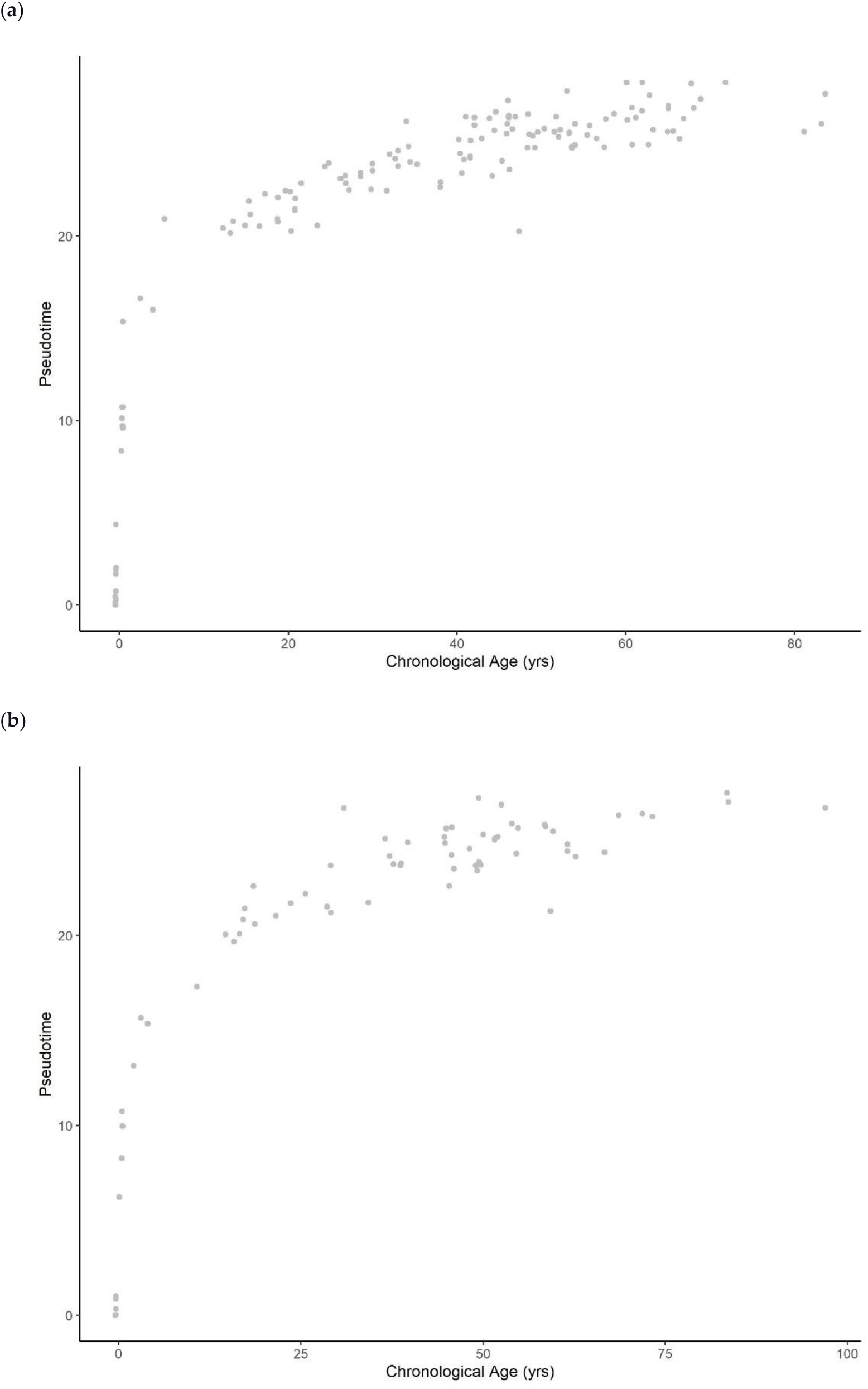
Methylation pseudotime of brain tissue samples in (**A**) train and (**B**) validation set.

**Figure A5.**
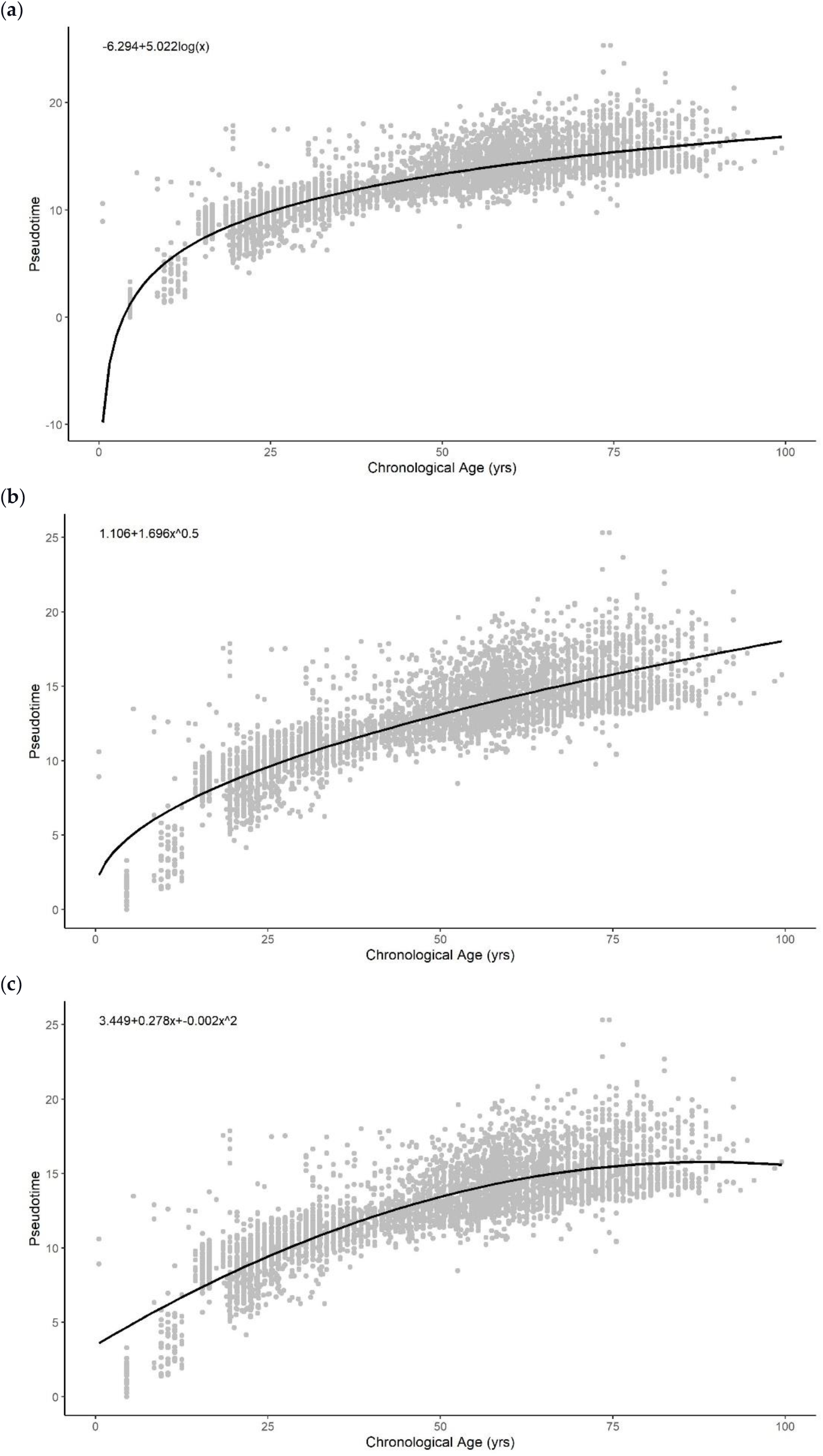
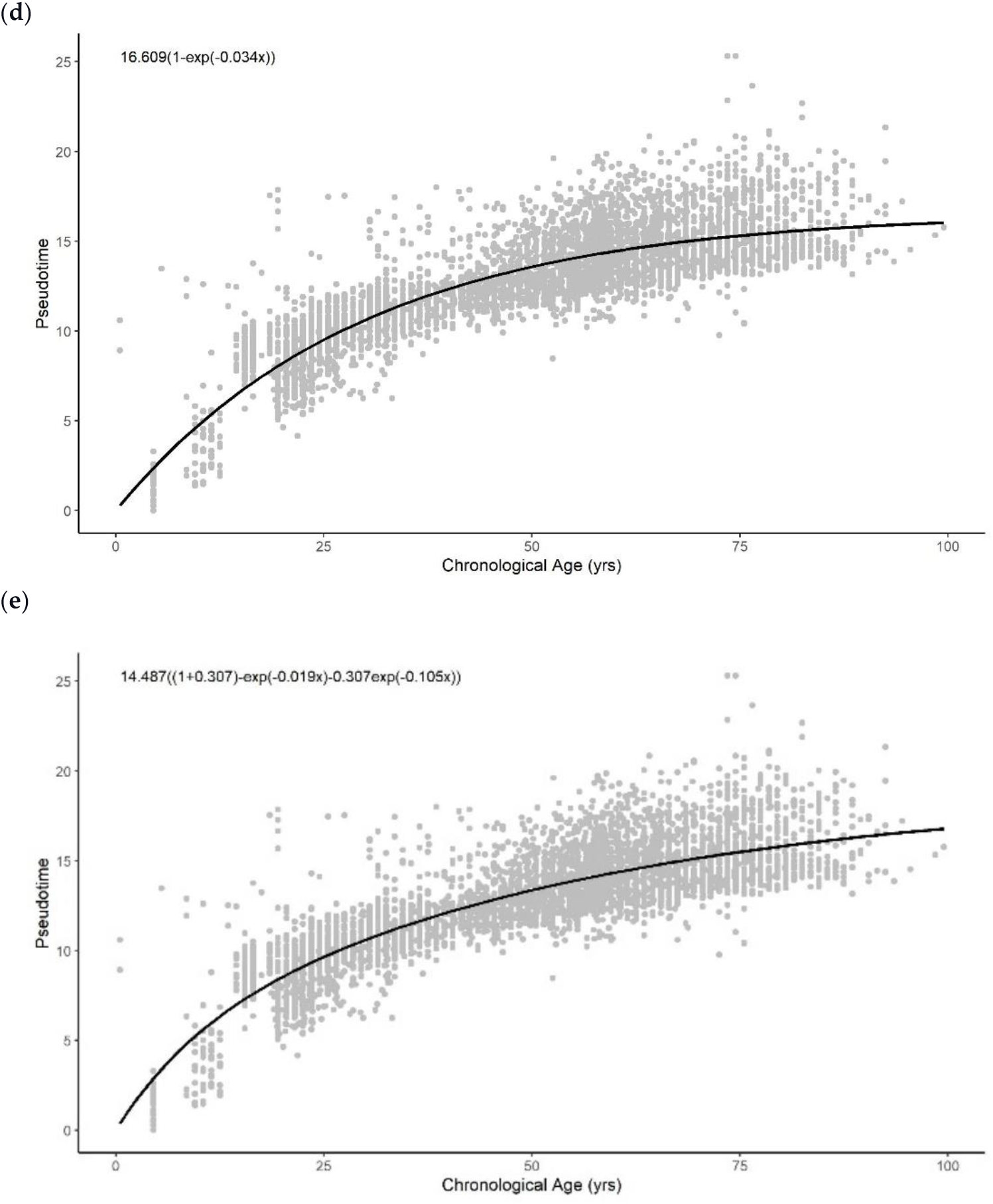
Methylation pseudotime of whole blood tissue samples with the best fitting sum of (**A**) logarithmic (**B**) square root (**C**) quadratic (**D**) exponential (**E**) sum of two exponential trendline

**Figure A6.**
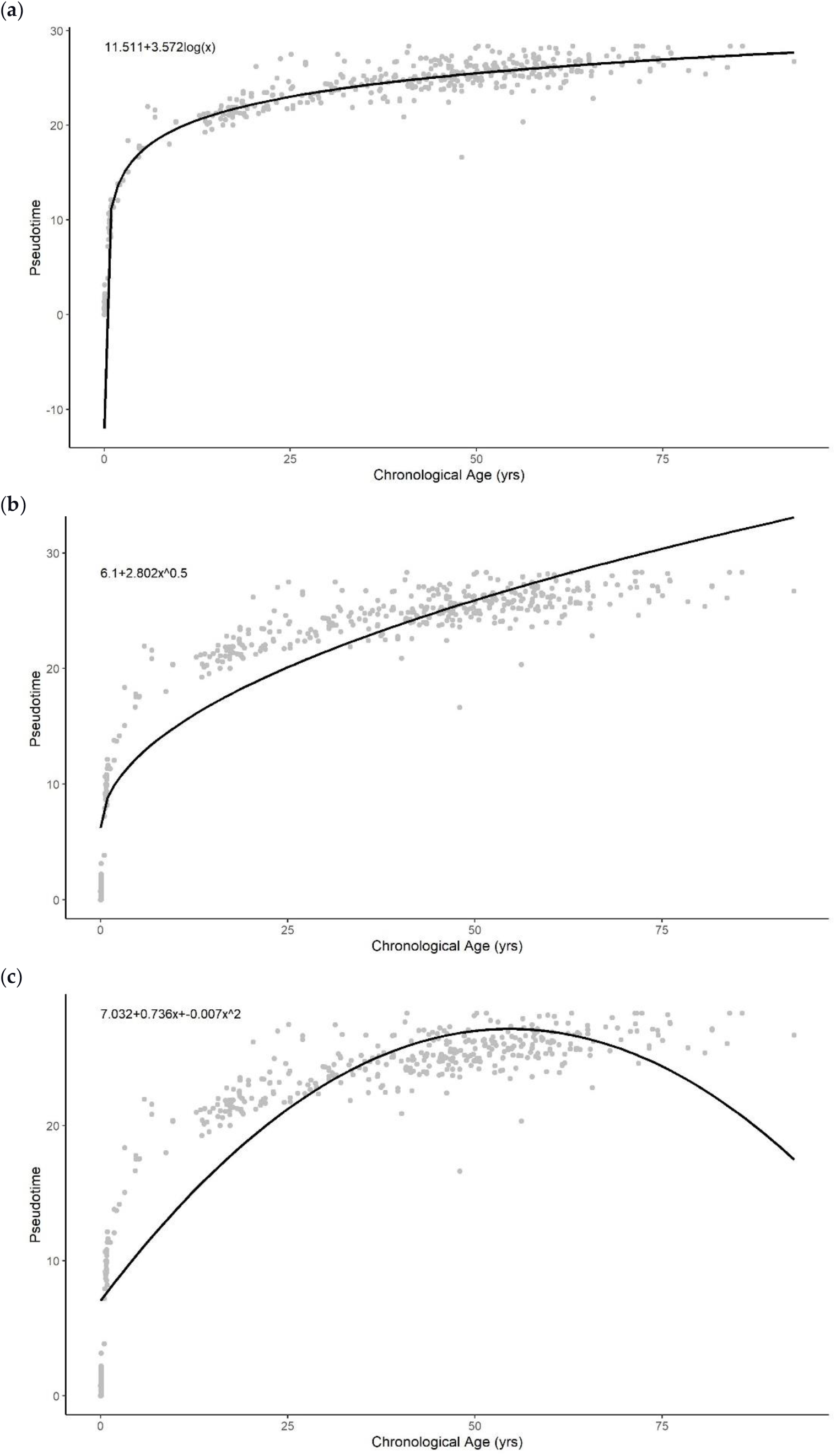
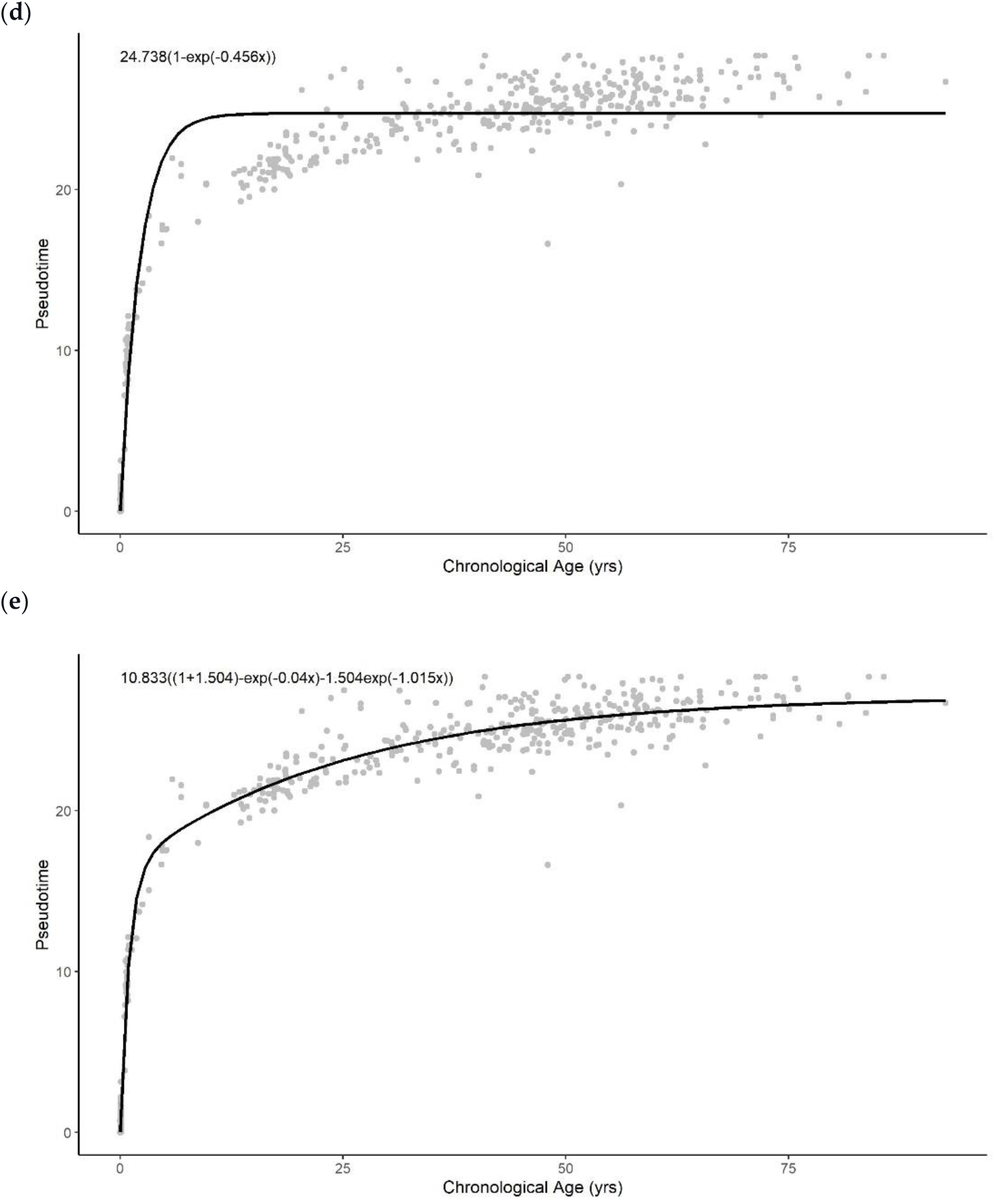
Methylation pseudotime of brain tissue samples with the best fitting sum of (**A**) logarithmic (**B**) square root (C) quadratic (**D**) exponential (**E**) sum of two exponential trendline

